# A scalable and modular computational pipeline for axonal connectomics: automated tracing and assembly of axons across serial sections

**DOI:** 10.1101/2024.06.11.598365

**Authors:** Russel Torres, Kevin Takasaki, Olga Gliko, Connor Laughland, Wan-Qing Yu, Emily Turschak, Ayana Hellevik, Pooja Balaram, Eric Perlman, William Silversmith, Forrest Collman, Uygar Sümbül, R. Clay Reid

**Author notes:** Equal contributors (Russel Torres, Kevin Takasaki, Olga Gliko).

## Abstract

Progress in histological methods and in microscope technology has enabled dense staining and imaging of axons over large brain volumes, but tracing axons over such volumes requires new computational tools for 3D reconstruction of data acquired from serial sections. We have developed a computational pipeline for automated tracing and volume assembly of densely stained axons imaged over serial sections, which leverages machine learning-based segmentation to enable stitching and alignment with the axon traces themselves. We validated this segmentation-driven approach to volume assembly and alignment of individual axons over centimeter-scale serial sections and show the application of the output traces for analysis of local orientation and for proofreading over aligned volumes. The pipeline is scalable, and combined with recent advances in experimental approaches, should enable new studies of mesoscale connectivity and function over the whole human brain.

## Introduction

Our knowledge of connectivity in the human brain is limited by the lack of high-resolution tools to study individual white-matter axons over long distances. Even the statistics of axon trajectories have not been studied with methods that can resolve individual axons. Studies of human brain organization have focused on mapping whole brains at low resolution, e.g. with diffusion MRI (Glasser et al. 2016; but see Walsh et al. 2021), or mapping small brain regions at extremely high resolution, e.g. with electron microscopy (Shapson-Coe et al. 2024), due to a lack of approaches that can both resolve individual axons and scale to the entire human brain. We have developed an approach, *axonal connectomics*, that builds on high-resolution, high-speed acquisition of fluorescently labeled *ex vivo* tissue samples using a combination of light-sheet fluorescence and tissue expansion microscopy (Turschak et al., 2026). At scale, axonal connectomics holds the promise of mapping both the local statistics and the long-distance targets of axon trajectories through white matter over entire human brains.

Despite significant advances in experimental methods (Ueda et al. 2020; Park et al. 2024), computational approaches to generating dense projection maps from the image data are needed. Analogous to serial section approaches in EM connectomics (Yin et al. 2020), the sample is first cut into sections in preparation for imaging, and subsequently, each section is imaged in a tiled acquisition from which the data must be montaged and reassembled into a volume *in silico* corresponding with the original sample. Furthermore, the problem of registering axons within a volume deformed by physical slicing, manipulation, and chemistry must be made solvable for datasets on the petavoxel scale.

We have developed a scalable computational pipeline for reconstructing and tracing axons for whole-brain axonal connectomics, building on progress in machine learning (ML)-based automated segmentation and tracing of axons (Gliko et al. 2024) and in petascale 3D registration and assembly of serially sectioned volumes (Mahalingam et al. 2022). Since our data consist almost entirely of trajectories of individual axons, rather than more general features typically seen in both light and electron microscopy, we used segmented skeletons to drive a novel pipeline for volume assembly. In contrast to previous approaches to volume assembly in electron and light microscopy utilizing computer vision techniques to extract salient corresponding features at or on the section interface and relying on local image statistics different from that of the axons themselves (Torres et al. 2020; Park et al. 2024), we investigated the use of the ML-segmented axonal traces as the basis for feature-detection and volume registration. Built on this foundation, we demonstrate a complete pipeline for axonal connectomics from data acquisition through proofreading and data analysis.

## Results

### Description of axonal connectomics pipeline

Tissue enters the data acquisition pipeline as sections which have been chemically fixed and immunostained for neurofilament proteins which densely label axons. These sections are highly scattering initially, but through a combination of tissue clearing and expansion steps adapted from standard expansion microscopy (ExM) protocols (Tillberg et al. 2016; Ku et al. 2020), proteins of interest are anchored to a hydrogel matrix while the surrounding tissue is digested and homogenized, rendering the sections optically clear (Fig. 1A). The hydrogel expands isotropically to 4x in water, enabling fluorescence imaging of axons with effective resolution better than the optical diffraction limit. Expanded tissue is imaged on a custom inverted selective plane illumination microscope (iSPIM) (Wu et al. 2011) in which the light-sheet and image plane are tilted such that the images are oblique cross-sections of the sample (Fig. 1B). By scanning the sample through the light-sheet while simultaneously imaging, the microscope acquires a 3D volume analogous to a z-stack in standard fluorescence microscopy, albeit with a skewed geometry. After imaging, the image tiles, or “strips,” are deskewed into Cartesian 3D volumes with nearly isotropic voxels of dimension 0.8 µm x 0.8 µm x 0.7 µm which corresponds to an effective resolution of 0.2 µm x 0.2 µm x 0.175 µm in unexpanded tissue. The deskewed tiles are stored in a next-generation file format, OME-Zarr (Moore et al. 2023), as multi-resolution pyramids with metadata recording the stage position and acquisition parameters used to place each tile volume in the coordinate system of the full section (Fig. 1C). This preprocessed data volume serves as the input to the tracing and assembly pipeline.

**Figure 1:**
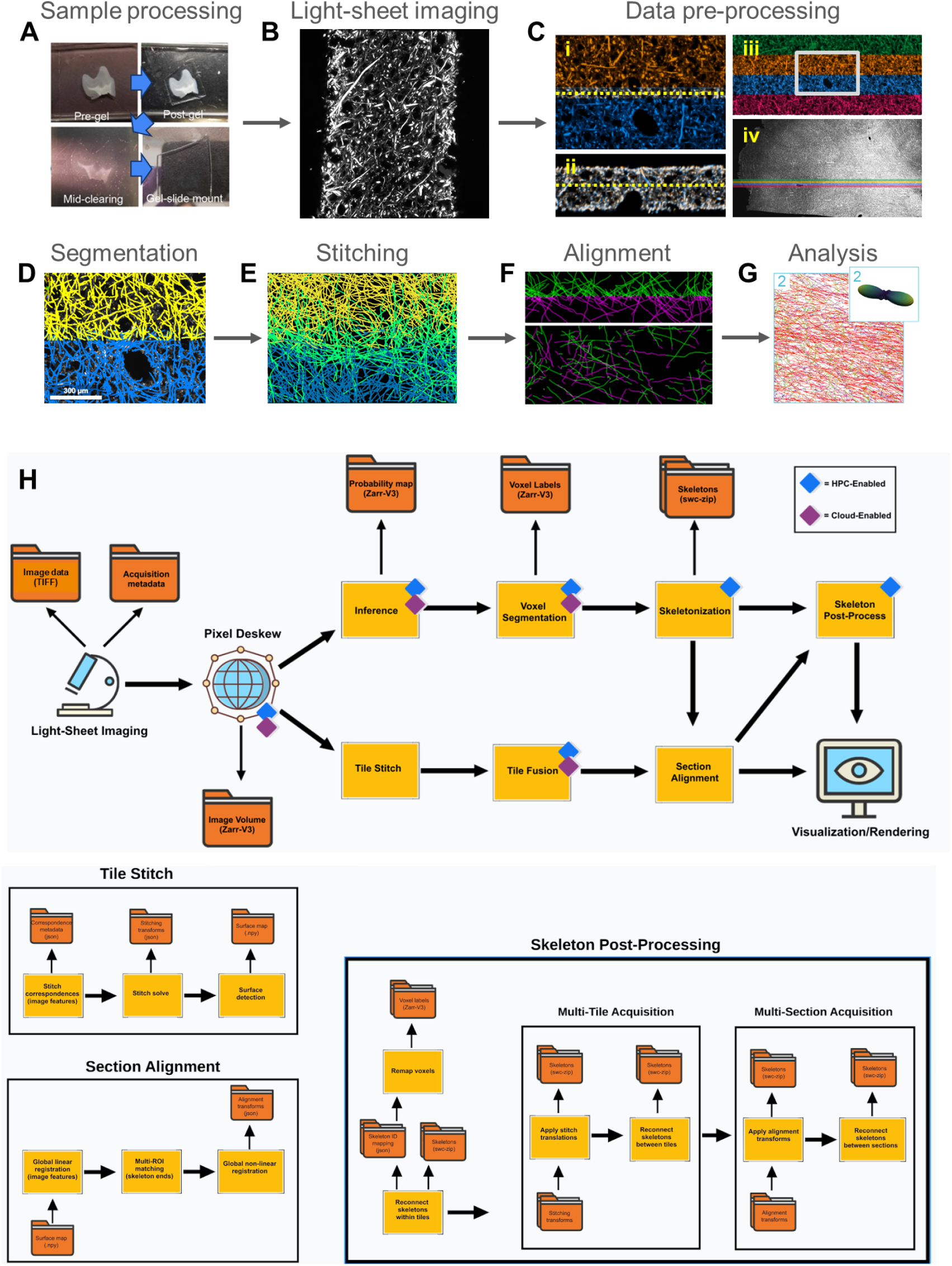
Pipeline for dense axon tracing over serially sectioned volumes. (**A**) Gelling, clearing, and expansion stages of tissue processing of immunostained sections. (**B**) Example fluorescence image from an expanded section acquired on an inverted light-sheet microscope. (**C**) Examples of adjacent, overlapping tiles (blue and yellow) after pre-processing image data into 3D volumes; (i): xy-plane sliced through dashed line in (ii); (ii): xz-plane sliced through dashed line in (i); (iii): set of 4 tiles with panel (i) shown in boxed region; (iv): zoomed out view of all section tiles with tiles in panel (iii) colored. (**D**) Example of axon traces segmented from blue and yellow tiles from (C). (**E**) Example of merged axons (green) stitched across blue and yellow tile boundaries. (**F**) Example of axon traces from two serial sections (green and magenta) aligned across their interface as viewed in the xz (top) and xy (bottom) planes. (**G**) Example of analysis of local axon orientation statistics. **(H)** Pipeline workflow defining the flow of image data from acquisition to visualization, and its accompanying intermediate outputs; (i): all modules aim to be agnostic to data location and compute resources, as shown by the progress made to enable high performance computing (HPC) and cloud paradigms; (ii): a more granular breakdown of the tile stitching, section alignment, and skeleton post-processing modules, all of which require the tracking of multiple intermediate outputs.

The computational pipeline encompasses a set of tools for segmentation, stitching, registration, transformation, and visualization of large image data, integrated into a modular workflow. Strips are first individually segmented, and the segmentation is skeletonized to generate traces of axons (Fig. 1D). Axon traces from adjacent strips are stitched in overlapping regions between tiles and merged across tile boundaries to register skeletons in a montaged section volume (Fig. 1E). Sections are aligned by matching and computing transformations to connect axon endpoints at surface interfaces (Fig. 1F). The outputs of the pipeline are the transformations needed for the registration of skeletons from disparate image tiles and sections into an assembled 3D volume, along with the sets of correspondences generating the transformations. The registered traces thus become available for proofreading and analysis (Fig. 1G).

Lastly, integration of the above tools in a workflow architected for cluster- and cloud-based computing is a design feature of the pipeline critical for deployment at scale. Here again, we adopted principles and tools developed for modern petascale EM pipelines and large-scale data infrastructure for managing distributed resources and processing (Fig. 1H).

### Automated segmentation and axon extraction

Our semi-automated (multi-step) approach for axonal reconstruction includes neural network-based segmentation followed by postprocessing and proofreading. The segmentation part of the computational pipeline uses OME-zarr image data of variable resolution as inputs and swc files as outputs, and allows generation of axon skeletons over entire sections. While proofreading is required for analysis of long-range axonal projections across the entire brain, it is not necessary for the section aligning process.

Axonal segmentation from 3D image stacks requires generating a voxel-wise annotation, like the one used for EM segmentation (Lee et al. 2021). Since it is a very compute-intensive task, we use a more time-efficient iterative approach by first segmenting dense axon image set, using an existing model trained with sparse neuron image sets (Gliko et al. 2024). We proofread automated skeletons and generate volumetric labels using a topology-preserved fast-marching algorithm (Gliko et al. 2024). We train a convolutional neural network (U-Net) to produce a voxel-based axonal segmentation. With this approach we are able to further refine the segmentation model with larger and more diverse datasets.

We algorithmically post-process voxel-based segmentation to obtain initial skeletons, which accurately capture axonal trajectories as assessed by human inspection, but contain topological errors such as false merges. The skeletons are therefore split at branch points, resulting in short linear segments, similar to the initial oversegmentation of EM data (Lee et al., 2021). We use the ground truth skeletons to train a classifier that learns a merging score, based on computed features for all pairs of nearby axonal segments. Segments are then iteratively merged, starting with the pairs with the highest score, reaching a conservative threshold chosen to minimize errors while increasing the mean length. This automated approach dramatically decreases the number of topological errors while reducing the proofreading time (Fig. 2A). We evaluate the quality of automated axonal reconstructions by comparing them to ground truth axons. Since only a limited part of axons was proofread, we select predicted axons with nodes that have corresponding nodes in ground truth within a distance of 2 voxels. We quantify the reconstruction accuracy using skeleton-based metrics as a fraction of ground truth axons being perfect, missing, truncated or split. We report these metrics for training and testing datasets (Fig. 2B). The number of splits per axon is in the 1-4 (1-8) range for testing (training) sets. We report an axonal segment as truncated if its length is below 90% of the length of ground truth axon and as split if it is both split and truncated. We do not report false merges due to incomplete ground truth, though we did not find any false merges by manual inspection of the test dataset. In addition, we found good correspondence between axon length, caliber and orientation for predicted and ground truth axons. Subsequent proofreading is performed to correct split errors and extend axon fragments into longer segments. Our proofreading on millimeter-scale regions of sections showed that >90% of axons can be traced to the edges of the volume, including the top and bottom surfaces of the section. This includes axons <1 µm in diameter, while a larger majority of the >1 µm axons are unbroken. This preliminary run-length analysis will form the backbone of our quality control metrics.

**Figure 2:**
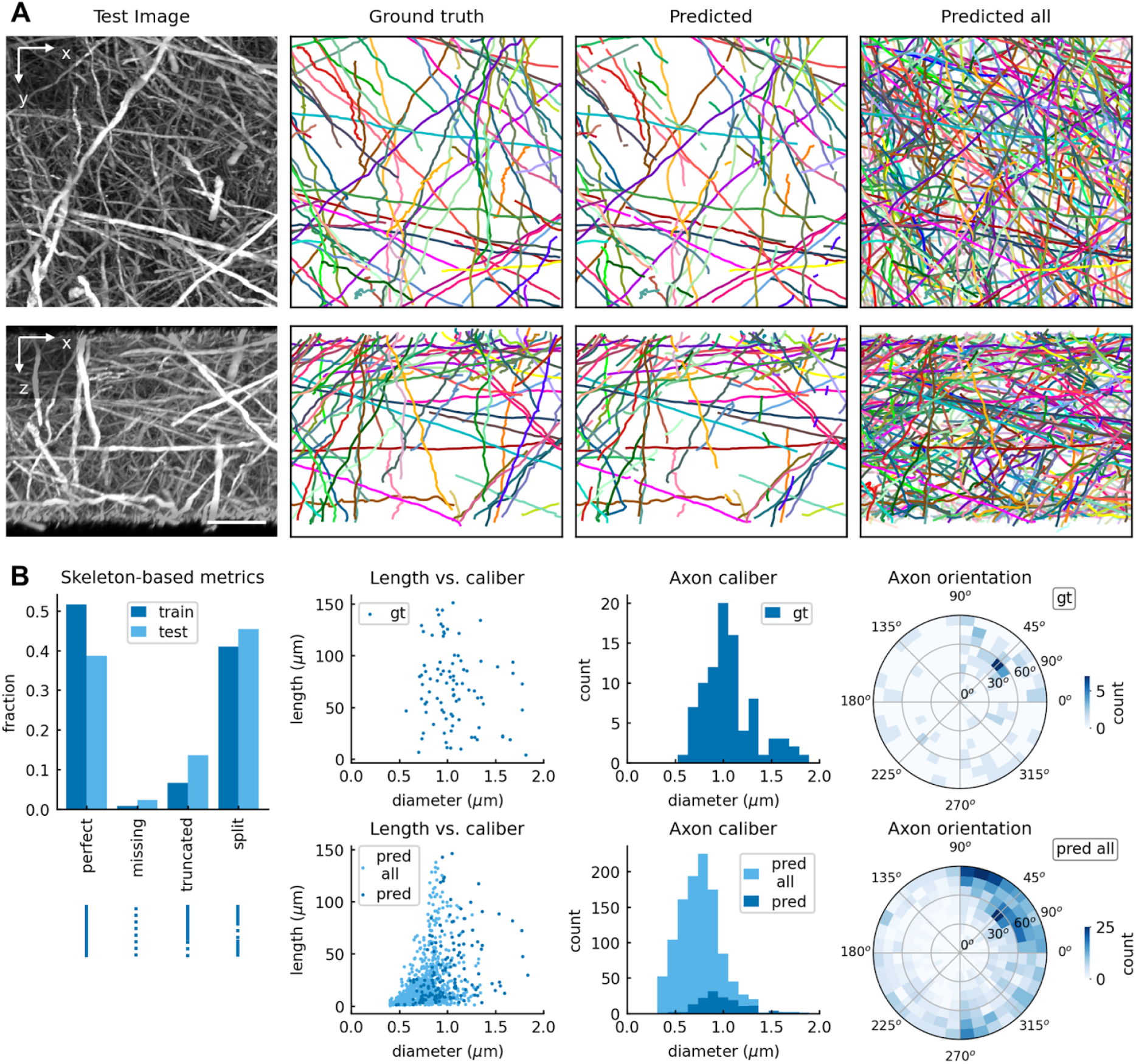
Automated axonal segmentation. (**A**) Segmentation of testing dataset: xy- and xz- maximum intensity projections of image stack (left column), corresponding plots of ground truth (gt) axons (second column), predicted axons matching gt (third column), and all predicted axons (right column). Different colors indicate different connected components (same color for the segments of split predicted axons matching gt). Scale bar, 25 µm. (**B**) Comparison of predicted and gt axons: skeleton-based metrics for training and testing dataset (left column), axon length vs. caliber for gt (second column, top) and predicted axons (bottom), caliber of gt (third column, top) and predicted axons (bottom), orientation (polar vs. azimuth angle heatmap) of gt (right column, top) and predicted axons (bottom).

### Stitching tiles into sections

Imaging of a large section is performed in tiles which must be co-registered to form a montage of the section in its entirety. For our image acquisition, a single section is tiled as a series of “strips,” each of which is generated by a unidirectional scan line along the x-axis of the stage and spaced across the y-axis such that ∼10% of the field-of-view overlaps between tiles for stitching (Fig. 3A). Due to inaccuracies of stage motion and other sources of uncertainty such as sample drift, the position of one tile in the imaging volume relative to another may deviate from the intended sample positioning. The stitching module generates and stores the output of stitching as metadata encompassing correspondences between spatial points in one tile with those of an adjacent neighbor and the set of transforms which register these point correspondences (Fig. 3B). These correspondences can be produced through a variety of standard image analysis techniques including correlation and template matching, as well as feature detection-based methods, but more generally, could be generated from more complicated representations, such as from neural networks and machine learning. Here, we use a multi-resolution combination of SIFT features (Lowe 1999) and cross-correlation for generation of stitching correspondences, and use colocalization of the ML-generated skeletons for parameter optimization and QC. The quality of stitching is commonly assessed from visualization and analysis of continuity of the transformed and rendered image data, and from identification of stitching-related artifacts, such as seams (Fig. 3C-D). The use of the skeletons for stitching validation enables automated QC and directly reports projection mapping errors.

**Figure 3:**
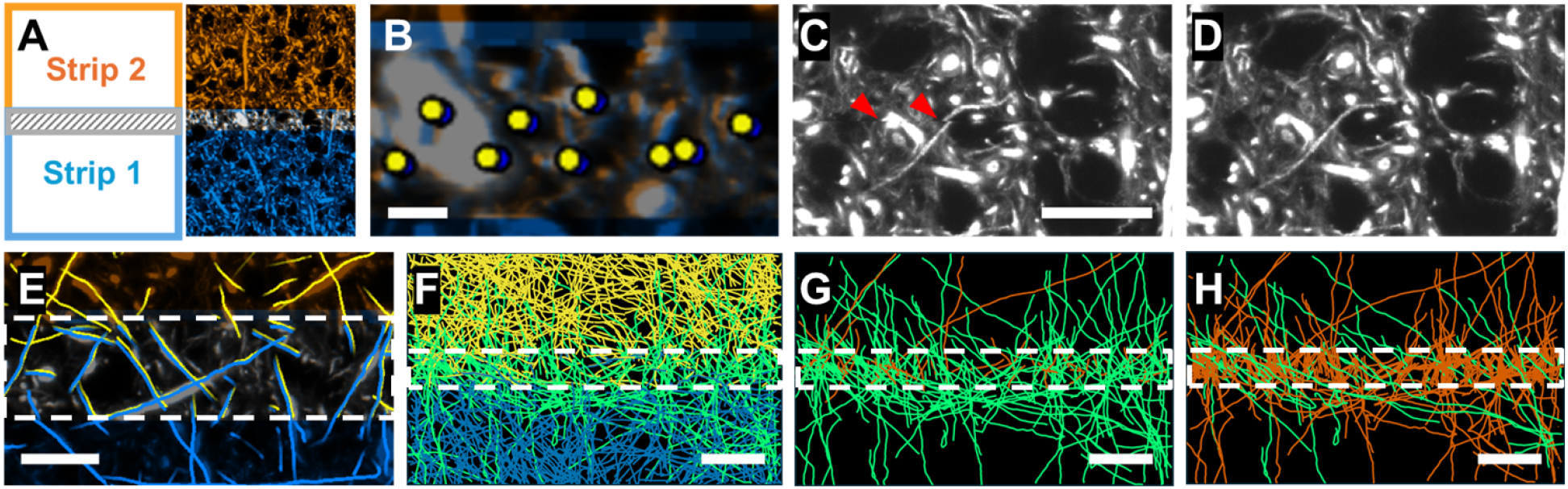
Automated stitching of axons within sections. (**A**) Schematic and example of overlapping strip acquisitions. (**B**) Stitching point correspondences generated in the overlap region. Scale bar, 20 µm. (**C**) Image data without stitching (stage position metadata only) at the boundary between tiles showing visibility of seams (arrows). Scale bar, 50 µm. (**D**) Fully stitched image data at the boundary in (C). (**E**) Stitched image data and skeletons in the overlap region between tiles (dashed lines). Scale bar, 50 µm. (**F**) Stitched skeletons merged across tile boundary (green) with skeletons from adjacent yellow and blue strips. (**G**) Merged skeletons (green) compared with unmerged skeletons present in the overlap region (red) from panel F. (**H**) Merged (green) and unmerged (red) skeletons in the overlap region from panel F without stitching. Scale bars, 100 µm.

Although minor misalignments may be visually subtle, they may still impact skeleton merging across tiles, which assumes near-complete overlap of the image data. Indeed, after transformation with the stitching translations determined from the image-derived correspondences, skeletons in the overlap region are expected to colocalize identically with the image data (Fig. 3E). Close alignment allows merging of the subset of skeletons that cross the tile boundaries by cutting skeletons at intersection points and joining the cut ends (Fig. 3F). Due to variability in the segmentation output between strips, as well as potential residual misalignment, a typically small fraction of skeletons present in the overlap region remain unmerged (Fig. 3G). Errors and failures of stitching are identifiable as a significant increase in the fraction of unmerged boundary skeletons (Fig. 3H), providing an intrinsic control on the quality of stitching.

### Assembling sections into volumes

The most challenging step in volume assembly is the unification of serial sections by tracing axons between slices. Our approach to alignment uses the endpoints of axon skeletons at the surfaces of adjacent sections, which are paired across the interface. Endpoint correspondences are generated using an iterative registration process between sub-regions of the dataset, flattened using a surface estimation, then jointly registered (Saalfeld et al. 2010; Khairy, Denisov, and Saalfeld 2018) using a progressively regularized set of 2D transformations as described in the ASAP volume assembly pipeline (Mahalingam et al. 2022).

First, highly downsampled image data (20 µm voxels) are used for the initial determination of a global rigid (rotation + translation) transform based on coarse registration of large anatomical landmarks such as major blood vessels (Fig. 4A). Following coarse registration and surface projection of endpoints over the entire section, the flattened 2D registration space is partitioned into a grid of subregions which can be individually aligned in parallel. Endpoints within each subregion are iteratively matched by running corresponding nearest neighbor, orientation angle, and intensity metrics through random sample consensus (RANSAC) regression, targeting a similarity to produce a stable set of inliers. These metrics isolate a subset of segmented axon endpoints for correspondence detection under the principle that the optimal transformation will result in the largest number of stable correspondences between filtered endpoints (Fig. 4B).

**Figure 4:**
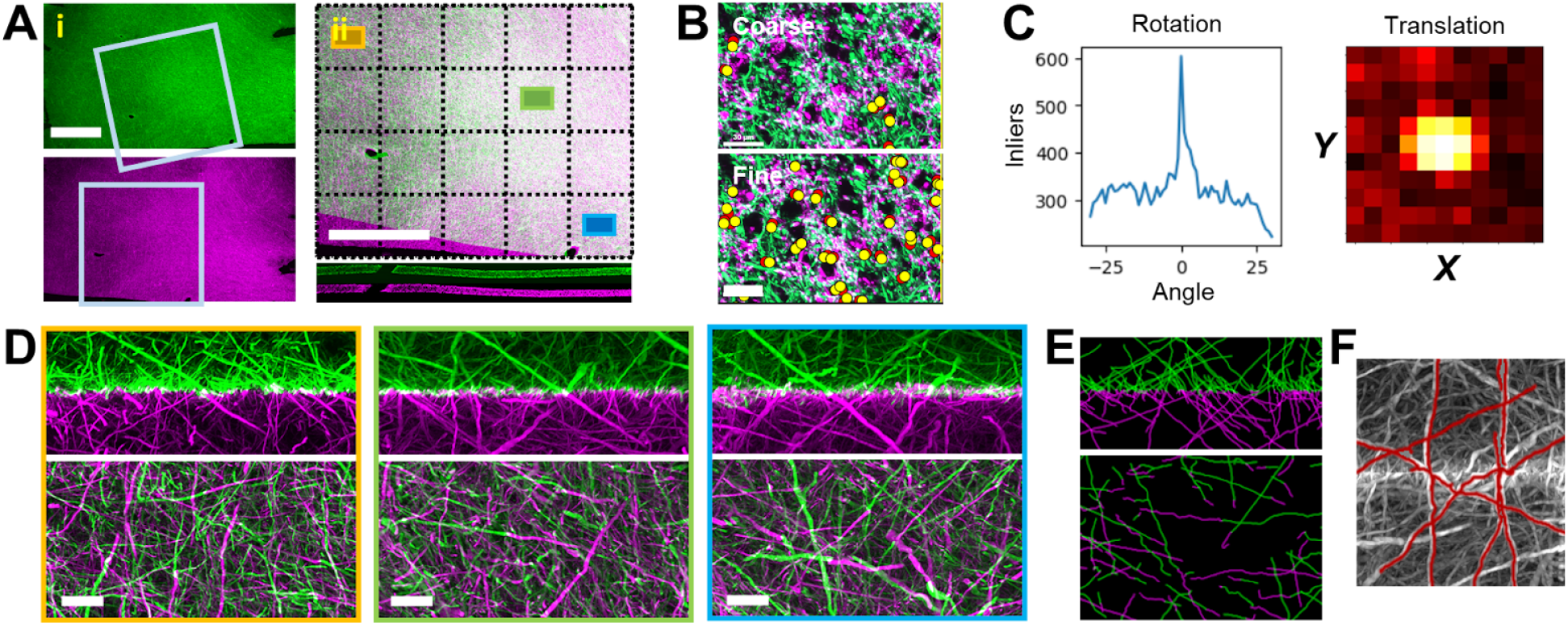
Automated registration and tracing of axons between serial sections. (**A**) Coarse registration of serial sections determined by matching large features (blood vessels); i: zoomed-out views of serial sections of human cortex (green, top, and magenta, bottom) with boxed field-of-view showing relative rotation; ii: zoom-in of coarse registration of boxed regions in panel (i) as viewed from the top (xy-slice, top) and from the side (xz-slice, bottom). Superimposed grid (dashed lines) shows partitioning into subregions for local correspondence detection and fine alignment. Scale bar, 5 mm. (**B**) Refinement of global (coarse, top) to local (fine, bottom) registration in subregions by similarity transformation optimization with consensus regression (RANSAC) over endpoint correspondences. Red and yellow markers indicate paired endpoints from top and bottom sections. Scale bar, 30 µm. (**C**) Fine alignment of endpoint correspondences by RANSAC and inlier optimization plotted against rotational (left) and translational (right) degrees of freedom. Peaks indicate maximum number of stable correspondences arising from locally optimized similarity transform. (**D**) Volume renderings of the transformed image data showing fine alignment of axons across the slice interface in boxed subregions outlined in yellow, green, and blue from aligned grid in A, scale bars: 100 µm. (**E**) Aligned skeletons from subregion as in D, matched, transformed, and 3D rendered at the flattened section interface. (**F**) Selection of skeletons crossing the section interface projected onto maximum intensity projection from flattened and aligned volume.

We generally observed good performance of similarity transformations within individual subregions for local optimization. Beginning with the global registration for the initial detection of correspondences with RANSAC, we solved for a local similarity transformation by iteratively optimizing the number of stable correspondences under rotation and translation (Fig. 4C). For subregions with filtered sets on the order of 1000 endpoints, this process typically completed in seconds on a single node and could be run in parallel on one computer or distributed across multiple nodes. For aggregating into larger or more deformed subregions, as well as a way to enforce consistency between subregions, an iterative approach to robust RANSAC allows greater coverage of correspondences while encompassing more local transformations (Saalfeld et al. 2012).The stable set of correspondence inliers across the section interface serve as input to a global alignment process solving across all section interfaces for increasingly regularized global rotational, affine, and grid-deformed thin-plate spline transformations. Transformation of the entire image data volume is not strictly necessary for the output of the pipeline given the connected traces are the primary output of section alignment. Further, transformations for very large regions can be generated by combining endpoint correspondences over sub-regions of interest, where transformations can be applied locally to for targeted rendering, validation, and proofreading (Fig. 4D-F and Supplementary Movies 1-2).

### Analysis of local orientation statistics and proofreading of axon traces

Following the complete computational pipeline for segmentation of densely labeled axons and image volume assembly, significant efforts in proofreading are required to trace individual axons by connecting pieces of split skeletons. Here, we present an exploratory analysis of local statistics of axon trajectory orientations using segmented skeleton data in the visual cortex of the human brain.

The max-z projection image of neurofilament immunostaining demonstrates densely labeled axon fibers interposed with each other (Fig. 5A). This region of interest was divided into 5x5 subregions, each covering approximately 500 x 500 µm in the xy plane. For each subregion, the orientation distribution function was estimated (Fig. 5B, left) based on the tangent vectors of skeleton segments within that region (Trinkle et al. 2021).

**Figure 5:**
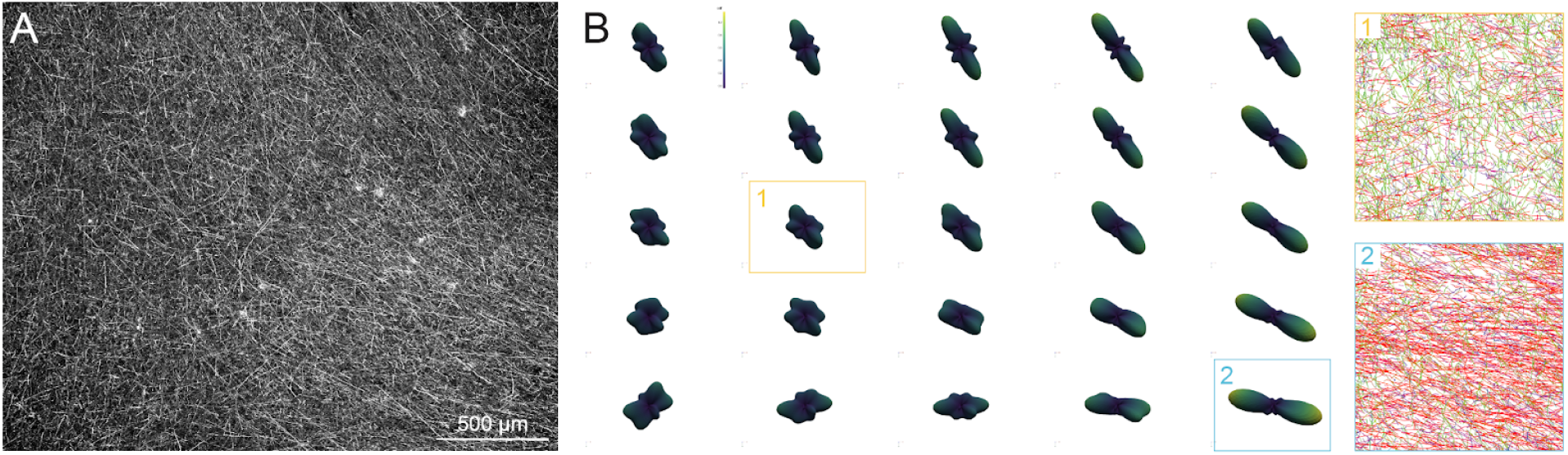
Analysis of local orientation of segmented axons. (**A**) Max-z projection view of image stacks from the visual cortex of human tissue covering about 2.5 x 2.5 x 0.12 mm pre-expansion, immunostained with neurofilament heavy-chain. Scale bar, 500 µm. (**B**) Left column: Orientation density function of segmented skeletons in each of 500 x 500 x 120 µm subregions. Zoomed-in views of skeletons in two selected regions are shown in the right column. Skeletons are colored by their orientation (orientation vector mapped to RGB space).

The analysis reveals a general global axonal trajectory from the lower right to the upper top. However, not all subregions exhibit a dominant axonal fiber orientation. In regions without a dominant fiber orientation (Fig. 5B, orange box), the orientation distribution is more uniform, indicating a more random arrangement of axonal trajectories. Even in regions with a dominant fiber orientation (Fig. 5B, blue box), there is still a significant number of axons that do not conform to the primary direction, highlighting the complexity and variability in axonal organization.

Finally, to enable scalable, interactive proofreading of the segmentation results, we developed a skeleton-guided supervoxel mapping and graph generation specifically adapted for ingestion into the ChunkedGraph format that serves as the entrypoint to the Connectome Annotation Versioning Engine (CAVE) infrastructure (Fig. 6) (Dorkenwald et al. 2022; Dorkenwald et al. 2025). While most CAVE datasets use derived supervoxels and graph edges from dense affinity-based EM segmentations, our methods leverage the topology of voxel-based skeletonization on a labeled segmentation to provide an entry point for skeletonized datasets from sparsely labeled data. We use Kimimaro (Silversmith et al. 2021) to oversegment the voxel labels, assigning labeled voxels to each proximal input skeleton vertex to generate new supervoxels (Fig. 6A-B). With this assignment, we derive the active supervoxel edges as well as the chunk-based components in the ChunkedGraph from the edges and vertices of the skeleton tree. To compensate for the difference in sparse labeling, we define inactive ChunkedGraph edges based on euclidean distance rather than supervoxel contact. This custom, skeleton-guided graph topology can be ingested into the hierarchical ChunkedGraph data structure same process as with EM segmentations to facilitate concurrent and efficient proofreading. As shown in Fig. 6C (also Supplementary Movie 3), when the automated segmentation produces a topological error such as a false split at the intersection of crossing axons, the established inactive graph edges allow users to perform merge operations within CAVE to reconnect the fragmented segments.

**Figure 6:**
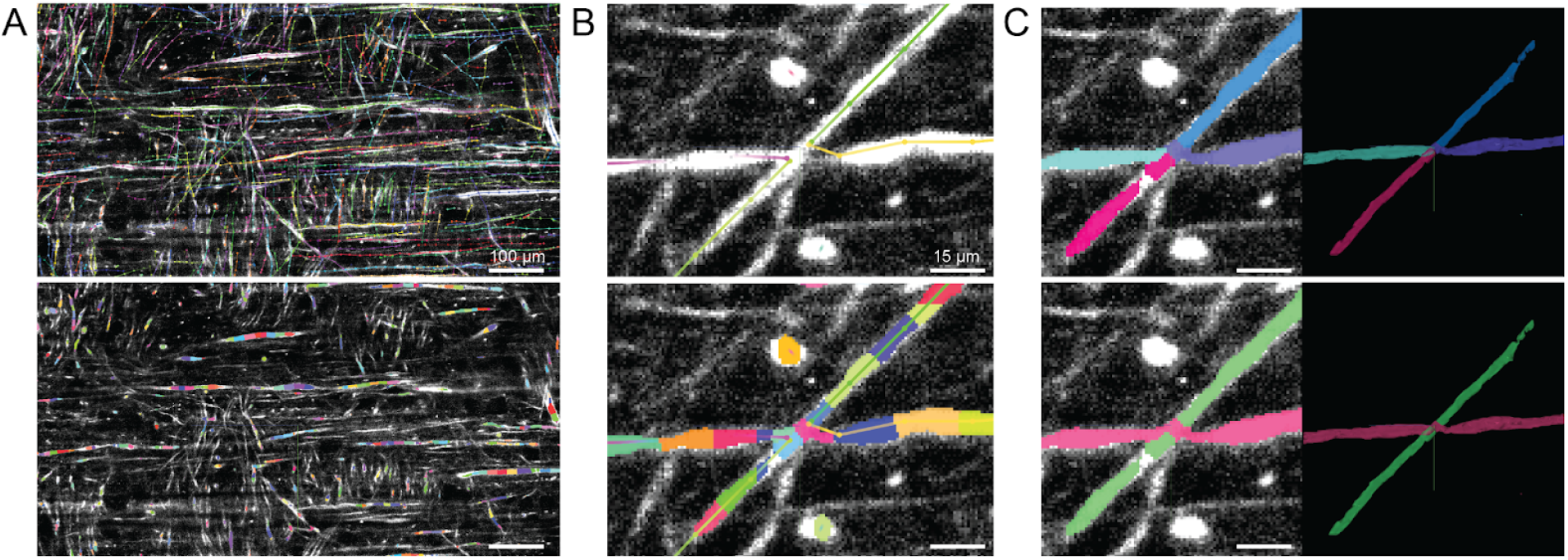
Skeleton-guided supervoxel mapping and interactive proofreading in dense axonal dataset. (**A**) Initial skeletons generated through the segmentation pipeline (top). The corresponding oversegmented supervoxels, where voxel labels are mapped directly to individual skeleton nodes (bottom). Scale bar, 100 µm. (**B**) Zoomed-in view of an intersection at a local crossing of two axons, whereas four skeleton fragments (top) directed the segmentation of image voxels into discrete supervoxels corresponding to each skeleton node. Graph edges connecting these atomic units are explicitly defined by the skeleton edges. (**C**) Demonstration of a merge operation to resolve this false split at the intersection by ingesting the skeleton-derived supervoxel graph into ChunkedGraph. The initial state before proofreading, showing a false split between two crossing axonal processes in the 2D view of segmentation (top left) and the 3D rendered mesh (top right). The corrected state after a user-guided merge edit, showing the same two correctly connected axons (bottom left and right). (See also Supplementary Movie 3). Scale bar, 15 µm.

## Discussion

### Comparisons with other approaches

The axonal connectomics tracing and assembly pipeline was designed to be a scalable and modular approach to generating dense projection maps from large, serially sectioned samples. The most similar alternatives to our knowledge are the serial sectioning and clearing, three-dimensional microscopy with semi-automated reconstruction and tracing (SMART) pipeline developed by the Bi group (Xu et al. 2021) and the Unification of Neighboring Sliced-tissues via Linkage of Interconnected Cut fiber Endpoints (UNSLICE) method developed by the Chung group (Park et al. 2024). The SMART approach in particular has demonstrated scalability to processing of entire primate brains; however, the pipeline was developed for comparatively sparse labeling of projections by viral injection and leverages sparsity in its alignment processing. The UNSLICE approach in contrast is able to align dense projections across serial sections but also requires co-staining of multiple molecular species and vasculature to perform step-wise registration in successive rounds of increasing accuracy with different imagery. An important distinction between these alternatives and the approach described here is the dense segmentation of axons providing high-quality traces for skeleton-based processing early in the pipeline and consequently, the primary role the segmentation plays in volume assembly. With the goal of projection mapping over large brains, we aimed to automate processing and validation with the traces as much as possible, and similarly minimize requirements for excess data acquisition and storage, as well as for processing of the raw image data. Given the difficulty and expense of acquiring, storing, managing, and processing extremely large data volumes, the scalability of our pipeline benefits from the economy of relying mainly on the processing of low-dimensional skeletons from only a single channel of data. Additional channels may still be acquired for non-alignment purposes, but each additional dataset comes at significant cost of increases in sample acquisition time and/or complexity, and computational and infrastructural burden.

Several published stitching software packages for large datasets are openly available (Hörl et al. 2019; Bria and Iannello 2012; Ruan et al. 2024). Existing stitching packages provide an integrated solution for both generating transformations and transforming the image data to render a “stitched” dataset. In our pipeline, the latter rendering step is a distinct process which we chose to avoid in many cases. Rendering stitched intermediates for processing was expensive and generally unnecessary given the existing segmentation and traces which we also aimed to leverage in the computation and validation of the stitching output. Thus, we implemented the lightweight stitching module described above with additional features for skeleton-based processing and QC. However, a key benefit of the modular design is that it is straightforward to substitute in the correspondences and transformations generated by another stitching package, which provides the opportunity for alternative implementations and further optimization in future development.

### Future directions

While our current segmentation captures the majority of axons and postprocessing allows us to correct a significant amount of false split/merge mistakes, much further improvement is needed to achieve more accurate tracing of axons. We expect to achieve significant improvement of segmentation accuracy by using larger and more diverse training data and employing affinity-based models with connectivity-preserving objective functions. In addition, our current approach for linking axonal segments obtained by splitting the branching nodes can be improved by training a graph neural network, where the nodes represent axonal segments and edge attributes capture relative position information, such as orientation and distance between a pair of segments.

The pipeline was built upon computational tools from the ASAP package developed for petascale volume assembly out of 2D image tiles from EM. In contrast, the image tiles acquired by the light-sheet microscope represent 3D volumes in themselves, and generally exhibited 3D deformations out of the plane which required supplementing the EM methods with modules for surface detection and flattening. Adaptations of EM tools were suitable for the projection mapping approach described here, namely the detection and registration of correspondences between axon traces across image tiles and section interfaces. However, the present pipeline does not support generalized volumetric transformations which becomes a limitation when aligning sections with severe surface curvature. We are currently investigating scalably extending our transformation models to handle 3D deformations.

We validated the proof-of-principle of segmentation-driven alignment by constructing the pipeline described above and applying it to axon tracing across a preliminary dataset of two serial sections. In a much larger effort combining multiple technical advances in sample preparation and imaging methods with the computational pipeline developed here, we are performing projection mapping over a volume at scale. As noted in the introduction, the problems of scaling include lifting constraints of sample preparation and of the field-of-view of imaging, which have been addressed by improvements in staining, clearing, and expanding thick sections and by recent innovation in light-sheet imaging technology (Glaser et al. 2023), respectively. The successful integration and demonstration of a fully scaled axonal connectomics pipeline promises to break new ground in our understanding of the large-scale structure of the human brain.

## Supplemental Methods

### Scalability and Distributed Processing

The pipeline is designed to scale to petavoxel-scale datasets by distributing computation across HPC resources using a combination of Nextflow, Gunpowder, and TensorStore. Most modules follow the same scalable processing paradigm. At the workflow level, Nextflow (a workflow framework for resource management) orchestrates parallel execution by allowing decomposition of large datasets into spatial subregions (ROIs) and dispatching them as independent jobs across HPC nodes. Within each job, the Gunpowder package provides a modular, streaming batch-processing framework that further subdivides each ROI into 3D chunks, which are read, processed, and written sequentially to an OME-Zarr dataset without loading the full volume into memory. Batch sizes are tuned per module and per resolution level to balance memory footprint against I/O overhead. Underlying all data access is the TensorStore package, which provides cloud and filesystem backed virtual array storage in chunked OME-Zarr format (in addition to most modern array storage filetypes). Because TensorStore supports concurrent reads and writes to non-overlapping regions, outputs from multiple parallel Nextflow processes can be written directly into a single consolidated output volume without requiring a separate merge step. Together, these tools allow the pipeline to process datasets spanning many teravoxels while remaining modular, fault-tolerant via retry logic, and straightforward to extend to new processing steps.

### Preprocessing: deskew, conversion, and multiscaling of acquisition data

Raw image data saved on the lightsheet microscope as TIFF stacks are uploaded to network storage and preprocessed by a dedicated server or cluster as described above. The file conversion process includes main data preprocessing routines for deskewing, downsampling, and compression. Deskewing of the image data to rectilinear coordinates generates voxels of 0.8 um x 0.8 um x 0.7 um dimensions at the highest resolution level which are then recursively 2x downsampled by block averaging to generate a 6-level *multum-in-parvo* (MIP) pyramid for multiresolution visualization and computation. The MIP level voxels are compressed with Blosc, generally achieving a 2-to 4-fold reduction in data size, and written to the chunked and sharded OME-zarr format.

### Equalization

Contrast equalization is applied during segmentation as a per-chunk pre-processing step to normalize intensity variations across large volumes. The pipeline uses percentile-based thresholding, whereby, for each chunk, intensities are rescaled by clipping to a low and high percentile, then linearly mapped to an 8bit range. This approach adapts to local intensity rather than applying a single global threshold, making it robust to the slow, large-scale intensity gradients common in expanded tissue. To avoid discontinuities at chunk boundaries, overlapping chunks are blended using a perimeter-weighted function. Each chunk is weighted by a spatially varying mask that smoothly tapers to zero at the edges, so that contributions from adjacent overlapping chunks blend seamlessly rather than producing sharp seams in the output.

### Segmentation model training

We iteratively refine our segmentation models using dense axon datasets. We generate automated segmentation using an existing model trained with sparse neuron image sets (Gliko et al. 2024). We use a neuron tracing interface, CATMAID (Saalfeld et al. 2009), to proofread automated skeletons and generate an initial set of ground truth skeletons. We use a topology-preserving fast marching algorithm to generate volumetric labels from proofread skeletons and image data (Gliko et al. 2024). Using these label and image sets as training data and employing data augmentation including flips/rotations by 90° in the image (XY) plane, brightness and contrast variations, we train a convolutional neural network (U-Net) to produce a voxel-based axonal segmentation. We use an efficient Pytorch implementation that runs on graphical processing units (GPUs) and train the model on batches of 3-d patches (128 x 128 x128 px^3^) with a GeForce GTX 1080 GPU using the Adam optimizer with a learning rate of 0.1.

### Skeletonization and Post-Processing

The initial voxel-based segmentation was postprocessed by thresholding the background, followed by connected component analysis to remove short segments, skeletonization with Kimimaro (Silversmith et al. 2021), conversion to digital reconstruction in swc format, and assigning a radius to every node.

Skeletonization, as all other modules ingesting array data, employs chunk based processing. As chunking naturally fragments contiguous skeletons, a conservative overlap is added to the bounding box of each chunk, and the resulting skeletons are merged together by identifying and merging matching vertices between chunks. Resolving fragmentation in this way allows us to scale HPC processing, whereby multiple regions of interest are processed concurrently with an overlap.

Post-processing includes breaking skeletons into fragments at branch points, optionally downsampling, and reconnecting fragments, all of which rely heavily on the Navis package of skeleton utilities. A classifier trained on ground truth data, using end distance and collinearity, is used to find putative matching pairs. These pairs are subsequently filtered by inference probability and merged to form longer axon traces. If multiple tiles and/or sections are processed, the resulting transforms are applied to relevant skeletons, and reconnection is run again.

### Within-section stitching

Stitching is performed on image data by extracting point correspondences in the overlapping region between adjacent tile pairs to calculate relative offsets of the tile volumes. Points of interest in the source tile are identified by spatial filtering and feature detection, such as with blob detection, and stage positioning metadata is used to determine “raw” correspondences, i.e. a set of coordinates in the destination tile corresponding to the set of source tile points according to the recorded stage motion. Cross-correlation is performed over limited 64 x 64 x 64 chunks around the raw correspondences at multiple resolutions to extract sub-voxel point correspondences between the source and destination tiles. Point correspondences between all pairs of tiles covering the section volume are submitted to a regularized least-squares solver to generate a set of tile offsets and translation-based montage representing the unified section.

### Between-section alignment

Alignment is performed after segmentation and skeletonization on the individual sections to be aligned and relies on detection of corresponding skeletons across the interface between sections. Section surfaces are detected using a series of image processing, filtering, thresholding, and edge detection steps at a low resolution, and represented as a gridded mesh. Skeleton endpoints near to the surface are extracted, and their positions and orientations, and endpoint intensities are calculated from the skeleton nodes and image voxels adjacent to the surface terminations, respectively. Surface endpoints covering 5000 x 5000 voxel matching subregions are identified based on an initial coarse rigid transformation and filtered for high angle orientations and high signal intensities. Optimization of endpoint pair alignment in a central 2000 x 2000 voxel region via a similarity transformation is performed with RANSAC and correspondence detection based on position, orientation, and intensity metrics with iterative searching of rotational and translational degrees of freedom. The optimized set of endpoint correspondences provides a local basis set for further fine alignment by nonlinear deformations with thin-plate splines, as well as aggregation of subregions by combining basis sets into larger areas of coverage. The correspondences identify skeletons of axons transiting the interface and enable tracing between sections.

### Volume Fusion

For materialization of overlapping tiles spanning an imaged volume, our pipeline provides a fusion module that places each tile into a common coordinate space using precomputed translation offsets, whether stage or registration derived, which specify the position of each tile’s origin within the global volume. Each tile is written into the appropriate location in a single output volume. If a surface map is provided, an optional surface-flattening step can be applied where each row of voxels along a chosen axis is shifted by a spatially varying offset, effectively unwarping curvature in the section before fusion.

### Proofreading with CAVE

The voxelized segmentation label data and corresponding skeletons, output by the processing pipeline, serve as input to a novel ingest strategy to a ChunkedGraph. We introduce a python module to facilitate integration of our data inputs with kimimaro (Silversmith et al., 2021) and pychunkedgraph (pcg). Our python library, ac_pcg, includes utilities to enable spatially chunked processing using kimimaro’s oversegment functionality, including efficient chunked spatial index based voxel relabeling with fastremap (Silversmith et al., 2025), as well as chunked spatial indexed filtering to map chunk edges and components based on the trees of pipeline-generated skeletons. For simplicity, this processing approach generates supervoxels that are entirely contained within spatial chunks of the dataset, meaning that the ChunkedGraph ingest consists entirely of within-chunk and between-chunk edges, without any cross-chunk edges.

ChunkedGraph initializations are also expected to be seeded with inactive edges, which we generated using a kdtree to find skeleton vertices outside of the querying connected component within a 30 voxel radius.

Ingest files for ChunkedGraph were generated for a subset of pipeline-generated data, using a jupyter notebook, and subsequently fed into a local deployment of the pychunkedgraph ingest service to produce a ChunkedGraph that could be used for proofreading.

Proofreading was accomplished using the tools provided by the CAVE infrastructure.

## Acknowledgments

We wish to thank the Allen Institute for Brain Science founder, Paul G Allen, for his vision, encouragement, and support. This work was supported by the National Institutes of Health through award numbers R01MH117820 and UG3MH126864 (PI: Reid), and UM1NS132207 (PI: Kamil Ugurbil, University of Minnesota).

## Author contributions

R.T., K.T., O.G., and C.R. conceived the pipeline. R.T., K.T., O.G., C.L., and W.Q.Y. contributed pipeline code, results, and analysis. E.T. prepared human histological sections. A.H. provided ground truth and proofreading. P.B., E.P., W.S., F.C., and U.S. provided methodology. All authors contributed to the manuscript.

## Literature cited

Bria, Alessandro, and Giulio Iannello. 2012. “TeraStitcher - a Tool for Fast Automatic 3D-Stitching of Teravoxel-Sized Microscopy Images.” BMC Bioinformatics 13 (November): 316.

Glaser, Adam, Jayaram Chandrashekar, Joshua Vasquez, Cameron Arshadi, Naveen Ouellette, Xiaoyun Jiang, Judith Baka, et al. 2023. “Expansion-Assisted Selective Plane Illumination Microscopy for Nanoscale Imaging of Centimeter-Scale Tissues.” ELife. 10.7554/elife.91979.1.

Glasser, Matthew F., Timothy S. Coalson, Emma C. Robinson, Carl D. Hacker, John Harwell, Essa Yacoub, Kamil Ugurbil, et al. 2016. “A Multi-Modal Parcellation of Human Cerebral Cortex.” Nature 536 (7615): 171–78.

Gliko, Olga, Matt Mallory, Rachel Dalley, Rohan Gala, James Gornet, Hongkui Zeng, Staci Sorensen, and Uygar Sümbül. 2024. “High-Throughput Analysis of Dendrite and Axonal Arbors Reveals Transcriptomic Correlates of Neuroanatomy.” Nature Communications 15:6337.

Hörl, David, Fabio Rojas Rusak, Friedrich Preusser, Paul Tillberg, Nadine Randel, Raghav K. Chhetri, Albert Cardona, et al. 2019. “BigStitcher: Reconstructing High-Resolution Image Datasets of Cleared and Expanded Samples.” Nature Methods 16 (9): 870–74.

Khairy, Khaled, Gennady Denisov, and Stephan Saalfeld. 2018. “Joint Deformable Registration of Large EM Image Volumes: A Matrix Solver Approach.” ArXiv [Cs.CV]. arXiv. http://arxiv.org/abs/1804.10019.

Ku, Taeyun, Webster Guan, Nicholas B. Evans, Chang Ho Sohn, Alexandre Albanese, Joon-Goon Kim, Matthew P. Frosch, and Kwanghun Chung. 2020. “Elasticizing Tissues for Reversible Shape Transformation and Accelerated Molecular Labeling.” Nature Methods 17 (6): 609–13.

Lee, Kisuk, Ran Lu, Kyle Luther, and H. Sebastian Seung. 2021. “Learning and Segmenting Dense Voxel Embeddings for 3D Neuron Reconstruction.” IEEE Transactions on Medical Imaging 40 (12): 3801–11.

Lowe, D. G. 1999. “Object Recognition from Local Scale-Invariant Features.” In Proceedings of the Seventh IEEE International Conference on Computer Vision, 2:1150–57 vol.2. IEEE.

Mahalingam, Gayathri, Russel Torres, Daniel Kapner, Eric T. Trautman, Tim Fliss, Shamishtaa Seshamani, Eric Perlman, et al. 2022. “A Scalable and Modular Automated Pipeline for Stitching of Large Electron Microscopy Datasets.” ELife 11 (July). 10.7554/eLife.76534.

Moore, Josh, Daniela Basurto-Lozada, Sébastien Besson, John Bogovic, Jordão Bragantini, Eva M. Brown, Jean-Marie Burel, et al. 2023. “OME-Zarr: A Cloud-Optimized Bioimaging File Format with International Community Support.” BioRxiv: 10.1101/2023.02.17.528834.

Park, Juhyuk, Ji Wang, Webster Guan, Lars A. Gjesteby, Dylan Pollack, Lee Kamentsky, Nicholas B. Evans, et al. 2024. “Integrated Platform for Multi-Scale Molecular Imaging and Phenotyping of the Human Brain.” Science 384,6701: eadh9979. doi:10.1126/science.adh9979

Ruan, Xiongtao, Matthew Mueller, Gaoxiang Liu, Frederik Görlitz, Tian-Ming Fu, Daniel E. Milkie, Joshua L. Lillvis, et al. 2024. “Image Processing Tools for Petabyte-Scale Light Sheet Microscopy Data.” Nature Methods 21, 2342–2352. doi:10.1038/s41592-024-02475-4

Saalfeld, Stephan, Albert Cardona, Volker Hartenstein, and Pavel Tomancak. 2009. “CATMAID: Collaborative Annotation Toolkit for Massive Amounts of Image Data.” Bioinformatics 25 (15): 1984–86.

Saalfeld, Stephan, Albert Cardona, Volker Hartenstein, and Pavel Tomančák. 2010. “As-Rigid-as-Possible Mosaicking and Serial Section Registration of Large SsTEM Datasets.” Bioinformatics 26 (12): i57–63.

Saalfeld, Stephan, Richard Fetter, Albert Cardona, and Pavel Tomancak. 2012. “Elastic Volume Reconstruction from Series of Ultra-Thin Microscopy Sections.” Nature Methods 9 (7): 717–20.

Shapson-Coe, Alexander, Michał Januszewski Daniel R. Berger, Art Pope, Yuelong Wu, Tim Blakely, Richard L. Schalek, et al. 2024. “A Petavoxel Fragment of Human Cerebral Cortex Reconstructed at Nanoscale Resolution.” Science 384 (6696): eadk4858.

Silversmith, William, J. Alexander Bae, Peter H. Li, A. M. Wilson. 2021. Seung-Lab/Kimimaro: Zenodo Release V1. 10.5281/zenodo.5539913.

Tillberg, Paul W., Fei Chen, Kiryl D. Piatkevich, Yongxin Zhao, Chih-Chieh Jay Yu, Brian P. English, Linyi Gao, et al. 2016. “Protein-Retention Expansion Microscopy of Cells and Tissues Labeled Using Standard Fluorescent Proteins and Antibodies.” Nature Biotechnology 34 (9): 987–92.

Trinkle, Scott, Sean Foxley, Narayanan Kasthuri, and Patrick La Rivière. 2021. “Synchrotron X-Ray Micro-CT as a Validation Dataset for Diffusion MRI in Whole Mouse Brain.” Magnetic Resonance in Medicine: Official Journal of the Society of Magnetic Resonance in Medicine / Society of Magnetic Resonance in Medicine 86 (2): 1067–76.

Turschak, Emily, Wan-Qing Yu, Kevin Takasaki, Steven J Cook, Russel Torres, Olga Gliko, Ayana Hellevik, Kareena Villalobos, Elizabeth Guadarrama, Soumya Chatterjee, Eric Perlman, Connor Laughland, Adam Glaser, Uygar Sümbül, R Clay Reid. 2026. “Heterogeneity of white-matter organization in the human brain.” BIORXIV/2026/714863.

Ueda, Hiroki R, Hans-Ulrich Dodt, Pavel Osten, Michael N Economo, Jayaram Chandrashekar, and Philipp J Keller. 2020. “Whole-Brain Profiling of Cells and Circuits in Mammals by Tissue Clearing and Light-Sheet Microscopy.” Neuron 106 (3): 369–387.

Walsh, C. L., P. Tafforeau, W. L. Wagner, D. J. Jafree, A. Bellier, C. Werlein, M. P. Kühnel, et al. 2021. “Imaging Intact Human Organs with Local Resolution of Cellular Structures Using Hierarchical Phase-Contrast Tomography.” Nature Methods 18 (12): 1532–41.

Wu, Yicong, Alireza Ghitani, Ryan Christensen, Anthony Santella, Zhuo Du, Gary Rondeau, Zhirong Bao, Daniel Colón-Ramos, and Hari Shroff. 2011. “Inverted Selective Plane Illumination Microscopy (iSPIM) Enables Coupled Cell Identity Lineaging and Neurodevelopmental Imaging in Caenorhabditis Elegans.” Proceedings of the National Academy of Sciences of the United States of America 108 (43): 17708–13.

Xu, Fang, Yan Shen, Lufeng Ding, Chao-Yu Yang, Heng Tan, Hao Wang, Qingyuan Zhu, et al. 2021. “High-Throughput Mapping of a Whole Rhesus Monkey Brain at Micrometer Resolution.” Nature Biotechnology 39 (12): 1521–28.

Yin, Wenjing, Derrick Brittain, Jay Borseth, Marie E. Scott, Derric Williams, Jedediah Perkins, Christopher S. Own, et al. 2020. “A Petascale Automated Imaging Pipeline for Mapping Neuronal Circuits with High-Throughput Transmission Electron Microscopy.” Nature Communications 11 (1): 4949.

